# The Clusters of Transcription Factors NFATC2, STAT5, GATA2, AP1, RUNX1 and EGR2 Binding Sites at the Induced *Il13* Enhancers Mediate *Il13* Gene Transcription in Response to Antigenic Stimulation

**DOI:** 10.1101/2020.09.29.318618

**Authors:** Mohammad Kamran, Jinyi Liang, Bing Liu, Yapeng Li, Junfeng Gao, Ashley Keating, Fathia Mohamed, Shaodong Dai, Richard Reinhardt, Yang Jiong, Zhongdao Wu, Hua Huang

**Affiliations:** Department of Immunology and Genomic Medicine, National Jewish Health, Denver, CO 80206; Department of Parasitology, Sun Yat-Sen University, Guangzhou, Guangdong 510800, China; Department of Immunology and Microbiology, University of Colorado School of Medicine, Denver, CO 80206; Department of Respiratory Medicine, Zhongnan Hospital of Wuhan University, Wuhan, Hubei 430071, China; University of Colorado Skaggs School of Pharmacy and Pharmaceutical Sciences

**Author notes:** These authors contribute equally (Mohammad Kamran, Jinyi Liang). Address correspondence and reprint requests to Dr. Hua Huang, Department of Biomedical Research, National Jewish Health and Department of Immunology and Microbiology, University of Colorado School of Medicine, 1400 Jackson Street, Denver, CO 80206.

## Abstract

Interleukin-13 plays a critical role in mediating many biological processes responsible for allergic inflammation. Mast cells express *Il13* mRNA and produce IL-13 protein in response to antigenic stimulation. Enhancers are essential in promoting gene transcription and are thought to activate transcription by delivering essential accessory co-factors to the promoter to potentiate gene transcription. However, enhancers mediating *Il13* have not been identified. Furthermore, which *Il13* enhancers detect signals triggered by antigenic stimulation have not yet been defined. In this study, we identified potential *Il13* enhancers using histone modification monomethylation at lysine residue 4 on histone 3 (H3K4me1) ChIP-seq and acetylation at lysine residue 27 on histone 3 (H3K27ac) ChIP-seq. We used Omni-ATAC-seq to determine which accessible regions within the potential *Il13* enhancers that responded to IgE receptor crosslinking. We also demonstrated that the transcription factor (TF) cluster consisting of the NFATC2, STAT5, GATA2, AP1, and RUNX1 binding sites at the proximal *Il13* enhancer, the TF cluster consisting of the EGR2-binding site at the distal *Il13* E+6.5 enhancer, are critical in sensing the signals triggered by antigenic stimulation. Those enhancers, which are responsive to antigenic stimulation and constitutively active, cooperate to generate greater transcriptional outputs. Our study reveals a novel mechanism underlying how antigenic stimulation induces robust *Il13* mRNA expression in mast cells.

## Introduction

The mouse interleukin-13 (*Il13*) gene contains 4 exons and 3 introns and is located at chromosome 11 (1). IL-13 is a pleiotropic cytokine produced by CD4^+^ Th2 T cells, CD8^+^ TC2 cells, natural killer (NK) T cells, basophils, and mast cells (2–7). It plays a critical role in the biological processes responsible for allergic inflammation (8). IL-13 induces goblet cell differentiation in the mucosal linings of the airway and the gut (9, 10), nitric oxide synthase in airway epithelium (11), fibroblast to myoblast transition (12, 13), and smooth muscle proliferation of airways (14). It also induces IgE class switching in B cells (15), leading to the production of antigen-specific IgE antibodies, which, together with their cognate antigens, activate basophils and mast cells.

Mast cells reside in the mucosal surface and connective tissues and express the high-affinity IgE receptor (FcεRI). They become activated when IgE and its cognate antigen bind and crosslink the FcεRI receptor (16, 17). Activated mast cells rapidly release stored inflammatory mediators, including histamine, heparin, and proteases. Mast cell activation also leads to *de novo* gene transcription of additional inflammatory mediators. Mast cells upregulate *Il13* gene expression and produce IL-13 protein in response to IgE receptor crosslinking (18) and also express the IL-33 receptor (ST) (19, 20). Stimulation of this receptor by IL-33, a cytokine produced by damaged epithelial cells, upregulates *Il13* gene expression, and production of the IL-13 protein (21–23).

Although several groups have investigated *Il13* gene regulation in T cells, enhancers that promote the *Il13* gene transcription in mast cells have not been studied systematically. In T cells, it has been reported that transcription factor (TF) GATA-3 is essential for the *Il13* gene transcription (24–27). GATA3 regulates *Il13* gene transcription by increasing chromatin accessibility at the *Il13* promoter to TF NFAT1 (28, 29). c-Myb promotes *Il13* gene transcription by interacting with GATA-3 at the GATA response element (30). In mast cells, NFAT and GATA2 cooperate to regulate *Il13* gene transcription (31). Masuda’s group described the interaction between GATA proteins and activator protein-AP1 at the promoter of *Il13* is necessary for *Il13* gene transcription in mast cells (32). Early growth response factor-1 (EGR1) is required for mast cells to transcribe *Il13* gene in response to IgE and antigen stimulation (33, 34). However, it is unknown which enhancers at the *Il13* gene respond to IgE and antigen stimulation.

TFs typically promote gene transcription by binding to clusters of TF-binding sites at enhancer regions. Binding of TFs to their corresponding binding sites at the enhancers leads to the recruitment of histone-modifying enzymes. The recruited histone methyltransferase MLL3/4 adds monomethylation at lysine residue 4 on histone 3 (H3K4me1), while the recruited histone acetyltransferase P300 acetylates lysine residue 27 on histone 3 (H3K27ac). Epigenomic studies demonstrate that H3K4me1 marks genes poised to be transcribed, whereas H3K27ac identifies genes that are actively transcribed (35–38). A newly developed technique, called Assay for Transposase-Accessible Chromatin (ATAC-seq), effectively identifies clusters of transcription factor binding sites at enhancers genome-wide (39, 40).

In this study, we used H3K4me1 and H3K27ac ChIP-seq to identify potential *Il13* enhancers, and Omni-ATAC-seq to identify clusters of TF binding sites at the potential *Il13* enhancers that responded to IgE receptor crosslinking. We constructed a series of *Il13* enhancer reporter genes to functionally verify clusters of TF binding sites that responded to IgE receptor crosslinking. We demonstrated that the TF cluster consisting of the NFATC2, STAT5, GATA2, AP1, and RUNX1 binding sites at the proximal *Il13* enhancer, the TF cluster consisting of the EGR2-binding sites at the distal *Il13* E+6.5 enhancer, are critical in sensing the signals triggered by antigenic stimulation. The IgE responsive and non-responsive enhancers cooperate to increase *Il13* gene transcription.

## Materials and Methods

### Cell culture

CFTL-15, a murine IL-3-dependent mast cell line (41), was provided by Dr. Melissa Brown (Northwestern University, Chicago, IL). CFTL-15 cells were grown in complete RPMI1640 medium supplemented with 10 % FBS, 100 units/mL penicillin, 100 μg/mL streptomycin, and 20 ng/mL IL-3 in 5% CO_2_ at 37°C Bone marrow-derived mast cells (BMMCs) were cultured from bone marrow cells of C57BL/6 mice in Iscove’s DMEM (10016CV, Corning TM cellgro TM, Manassas, VA) plus 10% FBS, 2 mM beta-mercaptoethanol 100 units/mL penicillin, 100 μg/mL streptomycin, and 20 ng/mL IL-3 for four weeks. Over 99% of BMMCs were mast cells, as determined by FACS analysis (FcεRI^+^ c-Kit^+^).

### Chromatin immunoprecipitation ChIP

Chromatin immunoprecipitation and ChIP-seq libraries were prepared as described before (42). Briefly, 1×10^7^ BMMCs were fixed by 1% formaldehyde. After sonication and pre-cleaning, the samples were incubated with either anti-H3K27ac antibody (ab4729, Abcam) or anti-H3K4me1 antibody (ab8895, Abcam) and then with protein A agarose/salmon sperm DNA slurry (Millipore, Cat# 16-157). The beads were washed and eluted according to protocols described before (42). The crosslinking of eluted immunocomplexes was reversed, and the recovered DNA was cleaned up using a QIAGEN QIAquick PCR purification kit (Qiagen, Valencia, CA). ChIP-seq data have been deposited in GEO (GSE145544).

### Omni-ATAC-seq

The Omni-ATAC-seq was performed according to the published method (40). Briefly, 50,000 BMMCs were either untreated or treated with DNP-IgE (1μg/mL, Sigma St Louis MO) for 12 hours followed by crosslinking with 10 μg/mL DNP-BSA (YAMASA, Tokyo, Japan) for 2 hours. The cells were spun down and washed with cold PBS once. Then the cells were resuspended in 50 μl cold ATAC-RSB-lysis buffer (10mM Tris-HCl pH 7.4, 10mM NaCl, 3mM MgCl2, 0.1% NP-40, 0.1% Tween-20 and 0.01% Digitonin) and incubated for 3 minutes. The lysis buffer was immediately washed out with 1 mL ATAC-RSB buffer (10mM Tris-HCl pH 7.4, 10mM NaCl, 3mM MgCl2, and 0.1% Tween-20). The cell pellets were resuspended in 50 μl transposition mix (25 μl 2X TD buffer, 2.5 μl transposase (Illumina, FC-121-1030), 16.5 μl PBS, 0.5 μl 1% digitonin, 0.5 μl 10% Tween-20, 5 μl H2O) and incubated at 37 °C for 30 minutes. The reaction was stopped by adding 2.5 μL of 0.5M EDTA pH 8, and transposed DNA was purified using the Qiagen MiniElute PCR purification kit (Qiagen). Purified DNA was amplified using the following condition: 72°C for 5 min, 98 *°C* for 30 s, and 7 cycles: 98 °C for 10 s, 63 °C for 30 s, 72 °C for 1 min. The amplified libraries were cleaned up, size-selected, and the quality and quantity of libraries were assessed on an Agilent Technologies 2100 Bioanalyzer. The pair-ended sequencing of DNA libraries was performed on an Illumina NovaSEQ6000 platform. ATAC-seq data has been submitted to GEO (GSE145542).

### Il13 promoter and enhancer constructions

The *Il13* minimal promoter (MP, −75 to −1 in relative to the transcription start site (TSS) of *Il13* gene) and P150 (−150 to −1) were cloned into pGL3-basic luciferase vector (Promega, Madison, WI) in BglII and HindIII cloning sites. The E-1.0 (−3208 to −1038 in the relative to TSS of *Il13* gene), E+0.2 (+238 to +1475), E+2.5 (+2533 to +4891), E+6.3 (+6534 to +8513), and E+8.9 (+8844 to +11372) enhancers were cloned into the DNA upstream of the *Il13* minimal promoter-luciferase vector through the NheI and BglII restriction sites. The combination of the *Il13* enhancers was generated using overlapping PCR as described (43) and cloned into the *Il13* minimal promoter-luciferase vector through the NheI and BglII restriction sites. Mutations in transcription factor binding sites were generated by using overlapping PCR as described (43) or using mutated oligos. The primers used for the amplification of enhancer fragments and mutated oligos are listed in Table S1. All mutations were verified by sequencing. All polymerases, and restriction and modification enzymes were obtained from New England Biolabs (Beverly, MA, USA).

### Transcription factor binding motif analysis

The putative transcription factor binding clusters were analyzed by using R/Bioconductor package TFBSTools (44). Transcription factor binding motif matrixes from the JASPAR 2016 database (http://jaspar.genereg.net/) were used in the analysis.

### Mast cell transfection, activation and enhancer reporter assay

CFTL-15 cells (1.25 ×10^6^) were electroporated with 10 μg luciferase plasmid and 0.5 μg Renilla vectors at 450 V and 500 μF using a Bio-Rad Gene Pulser. The transfected CFTL-15 mast cells were stimulated with DNP-IgE (1μg /mL) for 12 hours, followed by crosslinking with 10 μg/mL DNP-BSA for 6 hours (IgECL). Luciferase activities in the activated and control mast cells were measured with Infinite M1000^®^ microplate reader (Tecan Systems, Inc., San Jose, CA) using Dual-Luciferase reporter assay system (E1960, Promega). The transcription activity was measured as the ratio of luciferase and Renilla activity.

### Statistical analysis

All the error bars in this report represent mean ± standard deviation (mean ± SD). Statistical differences between the two samples were analyzed using the two-tailed student’s *t*-test.

## Results

### Identification of putative Il13 enhancers that respond to antigenic stimulation

We defined potential enhancers as noncoding DNA regions that are associated with both H3K4me1 and H3K27ac modifications. To identify potential *Il13* regulatory regions, we analyzed H3K4me1 ChIP-seq and H3K27ac ChIP-seq peaks at the *Il13* gene using the datasets generated and published by our laboratory (*Il13* enhancers were not analyzed in the paper, GSE97253). We showed that there were six potential enhancer regions within 14.6 kb of the mouse *Il13* gene, which covers the intergenic regions between the *Il4* and *Rad50* genes (Fig. 1A). We named these potential enhancers proximal enhancer (PE), E+8.9, E+6.5, E+2.5, E+0.2, and E-1.0 based on the distances of the putative enhancers to the transcription start site (TSS) of the *Il13* gene (Fig. 1A). Enhancers contain clusters of TF binding sites that can bind to TFs constitutively or bind to TFs activated by external stimulations. Binding of TFs to enhancers creates chromatin accessible regions within the enhancers (45). The chromatin accessible regions surrounding the TF bound sites can be identified by Assay for Transposase-Accessible Chromatin using sequencing (ATAC-seq). To assess which of the chromatin accessible regions can respond to IgE receptor crosslinking, we performed Omni-ATAC-seq, an improved version of ATAC-seq (40). We treated or did not treat bone marrow derived mast cells (BMMCs) with IgE receptor crosslinking for two hours. We observed that the chromatin accessible regions within the PE, E+8.9, E+6.5, E+2.5, E+0.2, and E-1.0 showed more accessibility in response to IgE receptor crosslinking (Fig. 1B). Sequencing data has been deposited to GEO (GSE145542). These results suggest that TFs are activated by IgE receptor crosslinking bound to the chromatin accessible regions at the PE, E+8.9, E+6.5, E+2.5, E+0.2, and E-1.0 enhancers either directly or indirectly through the transcriptional looping mechanism (45).

**Figure 1.**
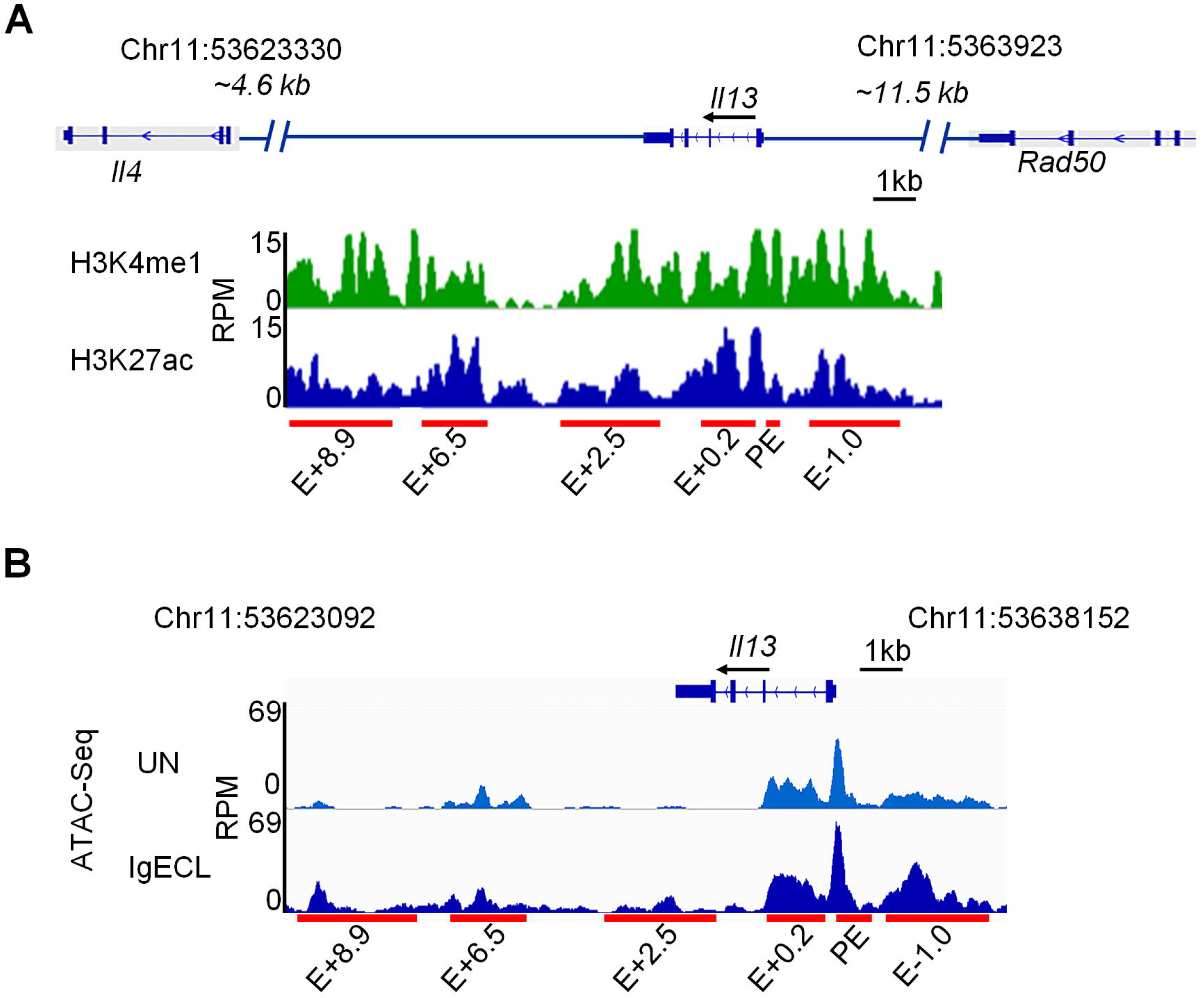
Identification of putative *Il13* enhancers. **(A)** H3K27ac and H3K4me1 ChIP-Seq tracks from chromosome position 53623092 to 53630153 (7,061 bp), which is located in the DNA regions between the *Il13* and *Il4* gene (4.6 kb to the TSS of the *Il4* gene) and between the *Il13* gene and Rad50 gene (11.5 kb to TSS of the *Il13* gene) in resting BMMCs. Red bars indicate putative *Il13* enhancers and promoter positions which showed significant H3K4me1 and H3K27ac modifications. E: enhancer; +0.2, +2.5, +6.3, +8.9 indicates the distance (kilo base pair) from the beginning of the downstream *Il13* enhancer to the TSS of the *Il13* gene, and −1.0 indicates the distance from the beginning of the upstream *Il13* enhancer to the TSS of the *Il13* gene. RPM: reads per million. **(B)** Omni-ATAC-seq tracks at the *Il13* gene obtained from BMMCs that were not stimulated (UN) or stimulated with IgE receptor crosslinking (IgECL). Data represent two biological samples.

Next, we used the luciferase reporter gene assay to determine which chromatin accessible regions at the six enhancers can respond to IgE receptor crosslinking directly. We constructed a series of enhancer reporter genes in which PE, E+8.9, E+6.5, E+2.5, E+0.2, and E-1.0 were linked to the *Il13* minimal promoter that contained only essential elements for RNA POL II binding (−75 to +1 relative to the transcription start site) (Fig. 2A). Mast cells cell-line CFTL-15 is a non-transformed, IL-3-dependent mast cell line (41). These cells are mature mast cells that respond to IgE receptor crosslinking with much higher cytokine gene transcription compared to transformed, immature, mast cell lines, such as the PT18 and MC/9 mouse mast cell lines. We transfected CFTL-15 mast cells with the series of constructs and treated or did not treat the transfected cells with IgE receptor crosslinking for 6 hours. We found that the PE had the highest induced enhancer activity in response to IgE receptor crosslinking (2.5-fold higher compared to the unstimulated group, Fig 2B). The E+6.5 enhancers also showed a moderate induced-enhancer activity after IgE receptor crosslinking (40% increase compared the unstimulated group, Fig. 2B), while the E-1.0 and E+0.2 possessed constitutively active enhancer activity with or without IgE receptor crosslinking. The E+2.5 and E+8.9 enhancers did not show significant enhancer activity with or without IgE receptor crosslinking (Fig. 2B). The results demonstrate that the clusters of TF binding sites at the accessible regions of PE and E+6.5 enhancer may bind to TFs activated by IgE receptor crosslinking.

**Figure 2.**
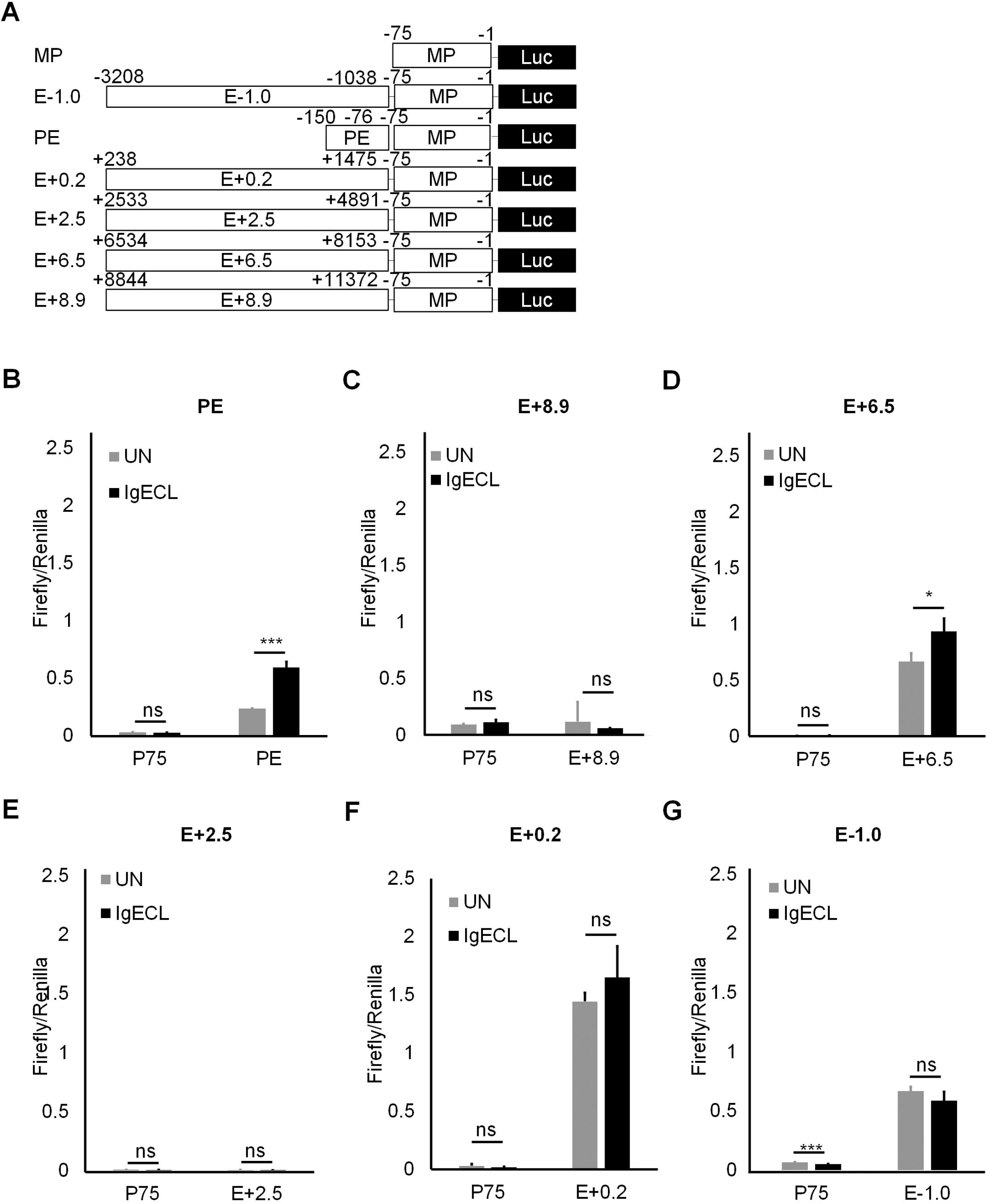
The PE and E+6.5 enhancer contain IgE-responsive elements. **(A)** Schematic diagrams showing enhancer reporter constructions. **(B)** Enhancer reporter analysis of PE enhancer transfected CFTL-15 cells that were not stimulated (UN) or stimulated with IgE receptor crosslinking (IgECL). **(C)** Enhancer reporter analysis of the E+8.9 enhancer. **(D)** Enhancer reporter analysis of the E+6.5 enhancer. **(E)** Enhancer reporter analysis of the E+2.5 enhancer. **(F**) Enhancer reporter analysis of the E+0.2 enhancer. **(G)** Enhancer reporter analysis of the E-0.1 enhancer. Mean ± SD were calculated from 3 transfectants in one experiment, representing two experiments with similar results. Statistical differences were analyzed by student’s t test. *: P<0.05, **: P<0.01, ***: P<0.001. MP, the minimal *Il13* promoter.

### Two inducible enhancers PE and E+6.5 cooperate with two constitutively active enhancers E-1.0 and E+0.2 to produce higher gene transcriptional outputs in response to IgE receptor crosslinking

Enhancers are thought to activate transcription by delivering necessary accessory co-factors to the promoter to potentiate gene transcription. Individual enhancers bound by TFs in the absence of IgE receptor crosslinking, and individual enhancers that bind to TFs activated by IgE receptor crosslinking, may loop to the promoter by the interaction of co-factors that bind to individual enhancers with the promoter. To determine whether the constitutively active enhancers and the IgE receptor crosslinking activated enhancers can act cooperatively and accumulatively, we constructed three enhancer reporter genes. In one, we linked all four enhancers that showed enhancer activity to the *Il13* minimal promoter. The second construct contained two inducible enhancers, and the third had two constitutively active enhancers (Fig. 3A). We observed that the constructs with two inducible enhancers and two constitutively active enhancers possessed the highest levels of transcriptional outputs under both resting and activated conditions (51-fold change in the resting and 86-fold in the activated condition compared to the minimal promoter). The construct that contained two inducible enhancers responded to IgE receptor crosslinking with increased gene transcription (28-fold increase in the resting, 54-fold in the activated CFTL-15 cells, respectively). The construct that contained two constitutively active enhancers did not respond to IgE receptor crosslinking. We concluded from these results that two inducible enhancers and two constitutively active enhancers cooperate to produce higher gene transcriptional outputs in response to IgE receptor crosslinking.

**Figure 3.**
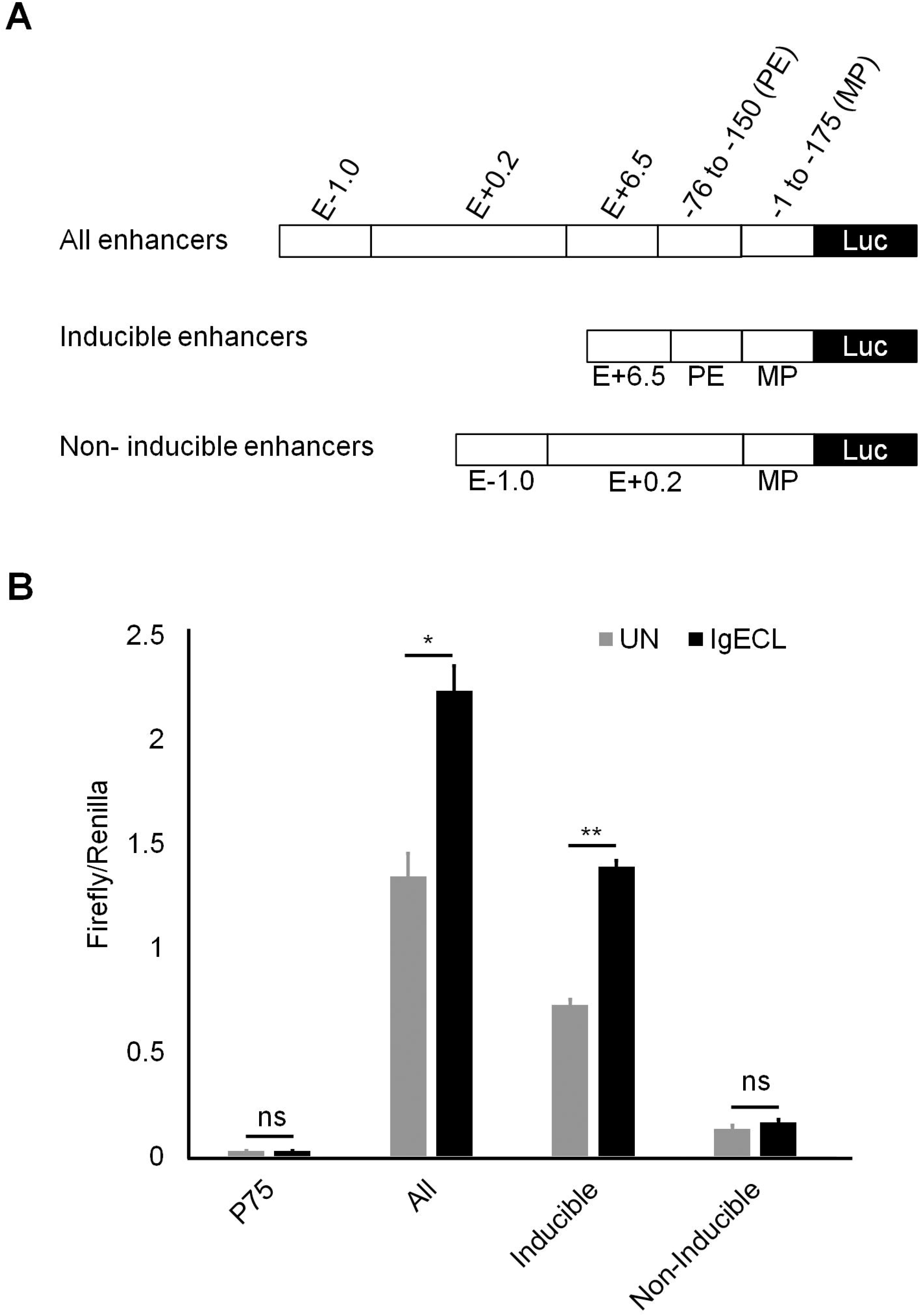
Two inducible enhancers PE and E+6.5 cooperate with two constitutively active enhancers E-1.0 and E+0.2 to generate higher gene transcriptional outputs. **(A)** Schematic diagrams showing enhancer reporter constructions. **(B)** Enhancer reporter analysis of the transfected CFTL-15 cells that were not stimulated (UN) stimulated with IgE receptor crosslinking (IgECL). Mean ± SD were calculated from 3 transfectants in one experiment, representing two experiments with similar results

### Combinatorial NFATC2, STAT5, GATA2, and AP1 binding sites at the PE enhancer are essential for the IgE receptor crosslinking-induced PE enhancer activity

To precisely determine which TF binding sites of the chromatin accessible regions are responsible for the IgE receptor crosslinking-induced PE enhancer activity, we performed TF binding motif analysis on the sequences found in the ATAC-seq peak in the PE enhancer (Fig. 4A). We found that consensus TFs NFATC2, STAT5, HOXA5, GATA2, AP1, and RUNX1 binding motifs at the region. To determine whether these TF binding motifs can promote the *Il13* reporter gene transcription, we performed mutational analysis of the TF binding sites. To reduce the numbers of permutations, we first generated a mutant PE enhancer, in which all potential TF binding sites were mutated. We then generated PE with the mutations at the overlapping TF binding sites and selected double or triple mutations that failed to respond to IgE receptor crosslinking for single TF binding site mutational analysis (Fig. 4A). As expected, mutation at all TF binding sites abolished the IgE receptor crosslinking-induced PE enhancer activity (Fig. 4B). We found mutations at both NFATC2 and STAT5 binding sites, but not single NFATC2 or STAT5 binding sites, abolished the IgE receptor crosslinking-induced PE enhancer activity (Fig. 4B). Similarly, triple mutations at GATA2, AP1 and RUNX1 binding sites eliminated the IgE receptor crosslinking-induced PE enhancer activity (Fig. 4B). Individual TF binding site mutation revealed that GATA2, AP1 and RUNX1 binding sites were critical in mediating the IgE receptor crosslinking-induced PE enhancer activity (Fig. 4B). In contrast, mutation at the HOXA5 binding site did not affect the IgE receptor crosslinking-induced PE enhancer activity (Fig. 4B). These results demonstrate that combinatorial NFATC2, STAT5, GATA2, AP1 and RUNX1 binding sites at the PE enhancer are required for the IgE receptor crosslinking-induced PE enhancer activity.

**Figure 4.**
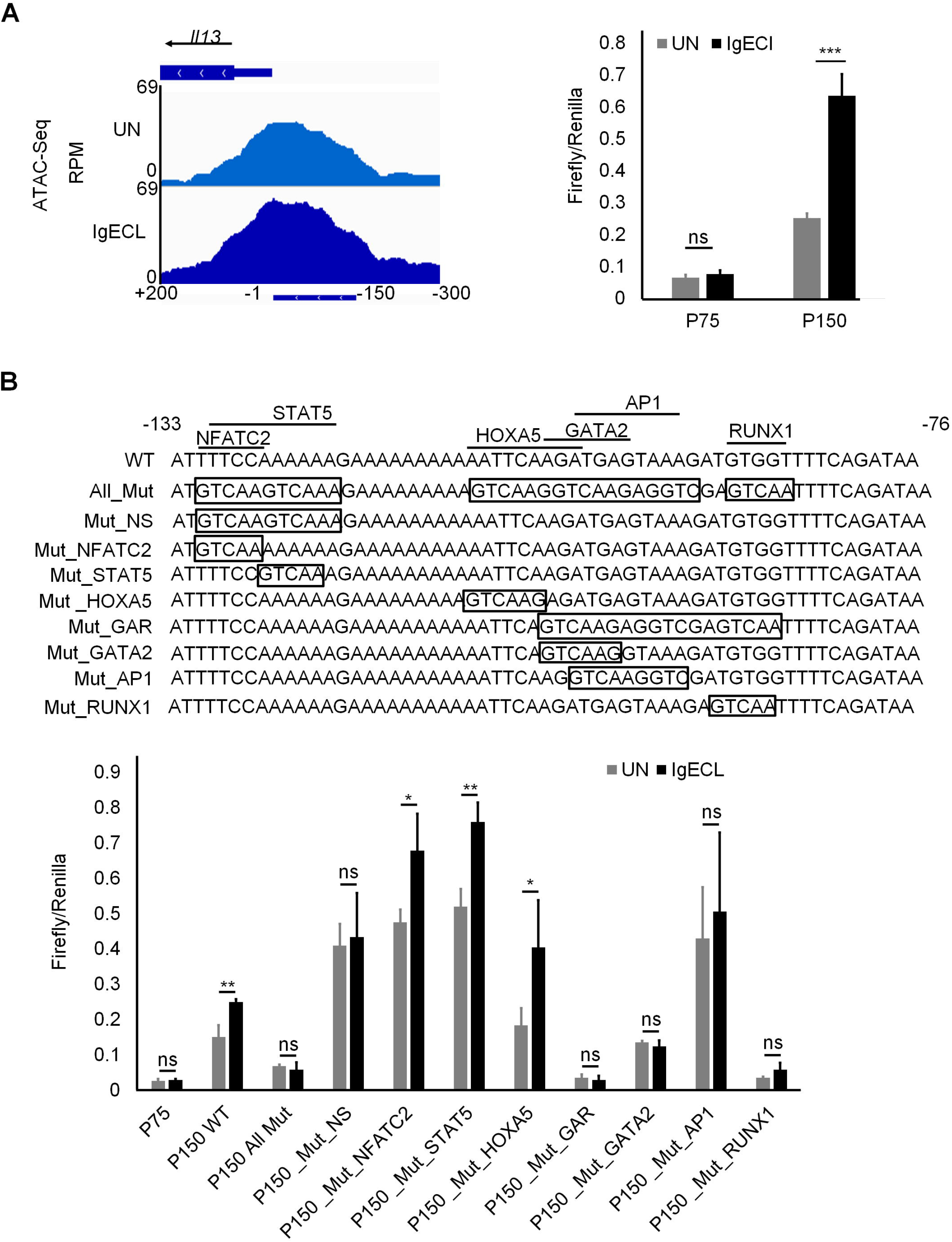
Combinatorial NFATC2, STAT5, GATA2, AP1 and RUNX1 binding sites at the PE enhancer are essential for the IgE receptor crosslinking-induced PE enhancer activity. **(A)** OmniATAC-seq tracks at the −300 to +200 region of the *Il13* gene in BMMCs that were not stimulated (UN) or stimulated with IgE receptor crosslinking (IgECL). Lower panel represents enhancer reporter analysis of the P150 enhancer reporter transfected CFTL-15 cells that were not stimulated (UN) stimulated with IgE receptor crosslinking (IgECL). **(B)** Upper panel depicts schematic diagrams showing various mutations at the PE region. Mutated sequences are indicated inside the boxed. Lower panel, enhancer reporter analysis of transfected CFTL-15 cells that were not stimulated (UN) stimulated with IgE receptor crosslinking (IgECL). Mean ± SD were calculated from 3 transfectants in one experiment, representing two experiments with similar results. Mut, mutant; Mut_NS, NFATC2 and STAT5 double mutants; Mut_GAR, GATA2, AP1 and RUNX1 triple mutants.

### The EGR2 binding site at the E+6.5 enhancer mediates the IgE receptor crosslinking-induced E+6.5 enhancer activity

To locate the IgE receptor crosslinking responsive TF binding sites within the E+6.5 enhancer, we first performed deletional analysis since the E+6.5 spanned over 2 kb long, making it challenging to conduct mutational analysis. We constructed a series of mutations by deleting 200 bp at a time (Fig. 5A, lower panel). We found that the deletion of a 200 bp DNA fragment from E+6.5_7370 to 7570, resulted in a loss of the IgE receptor-crosslinking-induced *Il13* reporter gene transcription, indicating this 200 bp DNA region contains clusters of TF binding sites that drove *Il13* reporter gene transcription in response to IgE receptor crosslinking. Indeed, this region was co-localized with an ATAC-seq peak (Fig. 5A), suggesting that this region is accessible to TFs. Further deletional analysis, in which we deleted DNA fragments 50 bp at a time, revealed that region between 7670 and 7720 was responsible for most of the *Il13* reporter gene transcription in response to IgE receptor crosslinking (Fig 5 B). TF binding analysis showed three potential TFs: FOXA2, EGR2, and STRE binding sites in the 50 bp fragment. We mutated FOXA2, EGR2, and STRE binding sites and found that the triple mutations at the 50 bp fragment abolished the IgE receptor crosslinking-induced *Il13* reporter gene transcription (Fig. 5C). Single mutation at the EGR2 binding site also dramatically reduced the IgE receptor crosslinking-induced *Il13* reporter gene transcription (Fig. 5C). Conversely, neither the mutation at the FOX2 or STRE binding site significantly decreased *Il13* reporter gene transcription in response to IgE receptor crosslinking (Fig. 5C). These results suggest that the EGR2 binding sites at the E+6.5 enhancer recruit TFs activated by IgE receptor crosslinking to promote *Il13* gene transcription in response to antigenic stimulation.

**Figure 5.**
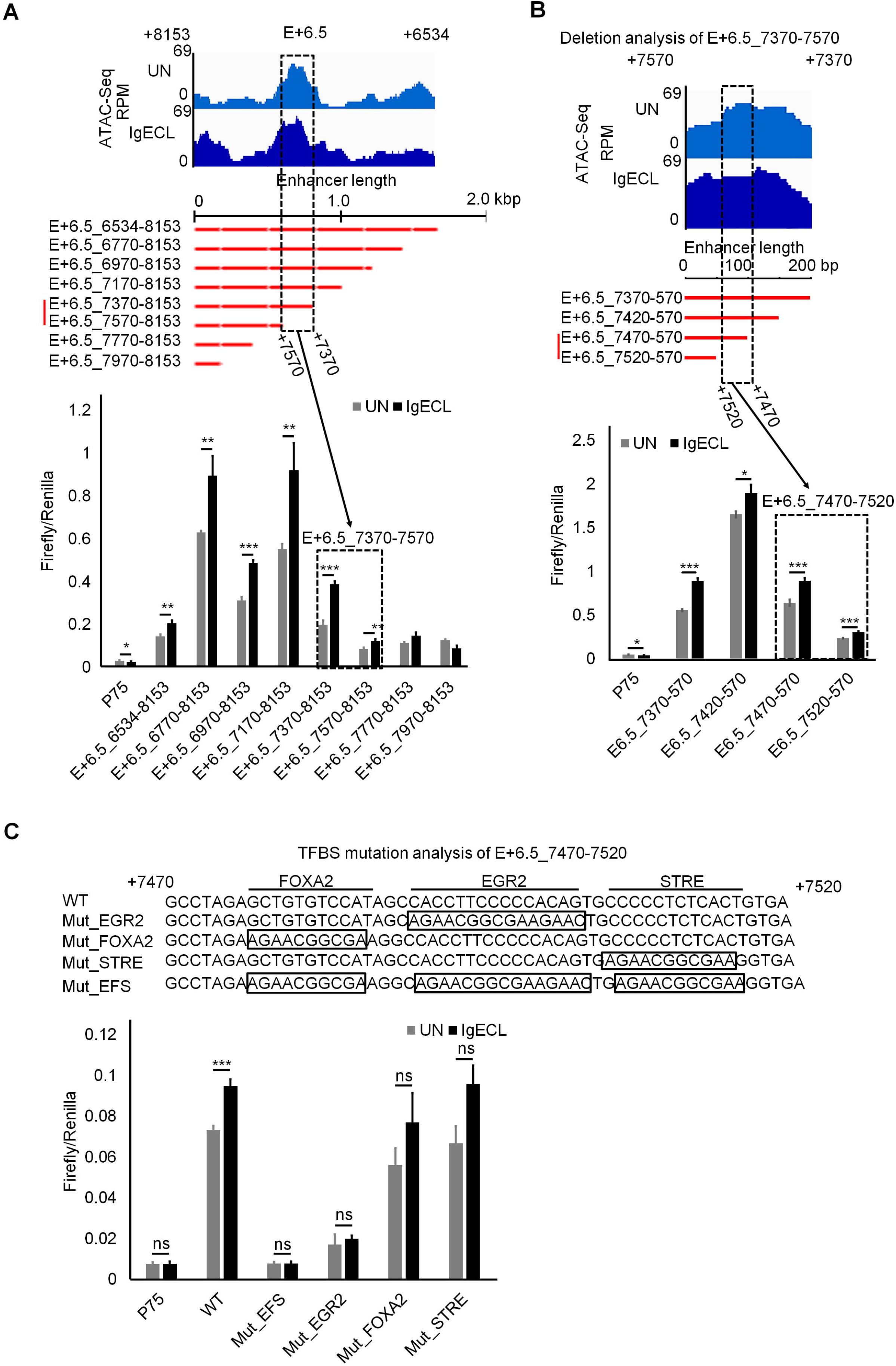
The EGR2 binding site at the E+6.5 enhancer mediates the IgE receptor crosslinking-induced E+6.5 enhancer activity. **(A)** Top panel shows Omni-ATAC-Seq peaks at the E+6.5 enhancers in resting (UN) and stimulated BMMC with IgE receptor crosslinking (IgECL). Middle panel is the schematic diagram of enhancer deletional analysis. The dotted box highlights the enhancer segments that lost its responsiveness to IgE receptor crosslinking. Lower panel, enhancer reporter analysis of the fragment from position +7370 to +7570 of the E+6.5 enhancer. **(B)** Enhancer reporter analysis of the fragment from position +7470 to +7520 of the E+6.5 enhancer. **(C)** Mutational analysis of the fragment from position +7470 to +7520 of the E+6.5 enhancer. Mut_EFS, EGR2, FOX2 and STRE triple mutants. Mean ± SD were calculated from 3 transfectants in one experiment, representing two experiments with similar results.

## Discussion

Systematic investigation of how *Il13* enhancers detect signals triggered by antigenic stimulation has not yet been explored in mast cells. In this study, we identified potential *Il13* enhancers using histone mark H3K4me1 and H3K27ac ChIP-seq. We then used Omni-ATAC-seq to determine which accessible regions within the potential *Il13* enhancers responded to IgE receptor crosslinking. We demonstrated that the TF cluster consisting of the NFATC2-, STAT5-, GATA2-, AP1- and RUNX1-binding sites in the *Il13* proximal promoter, the EGR2-binding sites in the *Il13* E+6.5 enhancer, respond to IgE receptor crosslinking as measured by their ability to direct *Il13* reporter gene transcription.

Our results support a model in that the clusters of TF binding sites at inducible PE and E+6.5 enhancers bind to activated TFs, and the binding recruits co-factors, such as mediators, that loop the distal E+6.5 to the *Il13* minimal promoter and PE enhancer. The formation of the E+6.5 enhancer, PE enhancer, and *Il13* promoter loop also bring together the constitutively active enhancers E-1.0 and E+0.2, which are located in the proximity of the *Il13* promoter and PE enhancers, presumably through the interaction of TFs and co-factors. We propose that the clusters of NFATC2-, STAT5-, GATA2-, AP1-, and RUNX1-binding sites at the PE bind to NFATC2, STAT5, GATA2, AP1, and RUNX1 either activated by IgE and antigen-mediated crosslinking of IgE receptors on the surface of mast cells or by cytokines produced by activated mast cells (in the case of STAT5) and that the EGR2-binding site at the inducible E+6.5 enhancer bind to EGR2 activated by IgE receptor crosslinking to induce the formation of transcription complexes in response to antigenic stimulation. Indeed, our previous study provides evidence that GATA2 plays a critical role in the *Il13* gene transcription in response to IgE receptor crosslinking (46). EGR2 was reported to be rapidly upregulated after IgE receptor crosslinking (47). It directly binds to the *Ccl1* chemokine gene promoter and is essential for *Ccl1*, but not for *Ccl3*, and *Ccl9* gene transcription and protein production (47). Thus, it is likely that EGR2 can bind to the EGR2-binding site at the E+6.5 enhancer to enhance the *Il13* gene transcription in response to antigenic stimulation.

A previous study showed that the DNA sequence (ttcaagatgagtaaa) located at the proximal *Il13* promoter from position −108 to −94 is a potential composite GATA2 and AP1 binding site (32). Single nucleotide mutation at the GATA2 and AP1-binding sites abolish luciferase reporter gene transcription in immature, transformed mast cell lines PT18 and MC9 mast cell lines that express low levels of *Il13* mRNA in response to IgE receptor crosslinking (31). Our results confirm the importance of the GATA2 and AP-1 composite sites in the *Il13* luciferase reporter gene transcription. Our results also demonstrate additional TFs NFATC2-, STAT5-, and RUNX1-binding sites in the PE enhancer that are important to mediating *Il13* luciferase reporter gene transcription. Using mature, IL-3-dependent, non-transformed mast cells CFTL-15 in our study might be critical in revealing this new finding. However, identifying the specific TFs that bind to the identified clusters of transcription factor-binding sites found in this study could be challenging. Because there are many potential TFs can bind to the same consensus TF binding site, future work that knock out or knock down individual TF genes in mast cells is needed to determine the exact TFs that bind to the consensus TF binding site at the enhancers we have identified.

In summary, IL-13 plays a critical role in mediating allergic inflammation, the underlying cause of many allergic diseases. Our detailed analysis of how TF binding sites at chromatin accessible regions at the IgE responsive enhancers in the *Il13* gene advances our understanding of mechanisms of allergic diseases.

## Acknowledgements

We thank lab members for thoughtful discussions. We are grateful to Ms. Marina Rahmani for technical assistance. We thank Ms. Kathleen Rauch for editor assistance.

## Correspondence address

Address correspondence and reprint requests to Dr. Hua Huang, Department of Immunology and Genomic Medicine, National Jewish Health and Department of Immunology and Microbiology, University of Colorado School of Medicine, 1400 Jackson Street, Denver, CO 80206. E-mail address

## Abbreviations

BMMCs: Bone Marrow-derived Mast Cells
IgE: Immunoglobulin E
TF: Transcription Factor
GATA2: GATA Binding Protein 2
GATA3: GATA Binding Protein 3
NFAT1: Nuclear Factor Of Activated T-Cells 1
AP1: Activator Protein 1
EGR1: Early Growth Response factor-1
STAT5: Signal Transducer and Activator of Transcription 5
RUNX1: Runt-related transcription factor 1
FOXA2: Forkhead box protein A2
NFATC2: Nuclear Factor Of Activated T Cells 2.

## References

1. McKenzie, A. N., X. Li, D. A. Largaespada, A. Sato, A. Kaneda, S. M. Zurawski, E. L. Doyle, A. Milatovich, U. Francke, and N. G. Copeland. 1993. Structural comparison and chromosomal localization of the human and mouse IL-13 genes. J Immunol 150: 5436–5444.

2. Brown, K. D., S. M. Zurawski, T. R. Mosmann, and G. Zurawski. 1989. A family of small inducible proteins secreted by leukocytes are members of a new superfamily that includes leukocyte and fibroblast-derived inflammatory agents, growth factors, and indicators of various activation processes. J Immunol 142: 679–687.

3. Cherwinski, H. M., J. H. Schumacher, K. D. Brown, and T. R. Mosmann. 1987. Two types of mouse helper T cell clone. III. Further differences in lymphokine synthesis between Th1 and Th2 clones revealed by RNA hybridization, functionally monospecific bioassays, and monoclonal antibodies. The Journal of experimental medicine 166: 1229–1244.

4. McKenzie, A. N., J. A. Culpepper, R. de Waal Malefyt, F. Briere, J. Punnonen, G. Aversa, A. Sato, W. Dang, B. G. Cocks, S. Menon, and et al. 1993. Interleukin 13, a T-cell-derived cytokine that regulates human monocyte and B-cell function. Proc Natl Acad Sci U S A 90: 3735–3739.

5. Burd, P. R., W. C. Thompson, E. E. Max, and F. C. Mills. 1995. Activated mast cells produce interleukin 13. The Journal of experimental medicine 181: 1373–1380.

6. Akbari, O., P. Stock, E. Meyer, M. Kronenberg, S. Sidobre, T. Nakayama, M. Taniguchi, M. J. Grusby, R. H. DeKruyff, and D. T. Umetsu. 2003. Essential role of NKT cells producing IL-4 and IL-13 in the development of allergen-induced airway hyperreactivity. Nature medicine 9: 582–588.

7. Ochensberger, B., G. C. Daepp, S. Rihs, and C. A. Dahinden. 1996. Human blood basophils produce interleukin-13 in response to IgE-receptor-dependent and - independent activation. Blood 88: 3028–3037.

8. Wills-Karp, M., J. Luyimbazi, X. Xu, B. Schofield, T. Y. Neben, C. L. Karp, and D. D. Donaldson. 1998. Interleukin-13: central mediator of allergic asthma. Science 282: 2258–2261.

9. Kanoh, S., T. Tanabe, and B. K. Rubin. 2011. IL-13-induced MUC5AC production and goblet cell differentiation is steroid resistant in human airway cells. Clin Exp Allergy 41: 1747–1756.

10. Khan, W. I., P. Blennerhasset, C. Ma, K. I. Matthaei, and S. M. Collins. 2001. Stat6 dependent goblet cell hyperplasia during intestinal nematode infection. Parasite immunology 23: 39–42.

11. Suresh, V., J. D. Mih, and S. C. George. 2007. Measurement of IL-13-induced iNOS-derived gas phase nitric oxide in human bronchial epithelial cells. Am J Respir Cell Mol Biol 37: 97–104.

12. Saito, A., H. Okazaki, I. Sugawara, K. Yamamoto, and H. Takizawa. 2003. Potential action of IL-4 and IL-13 as fibrogenic factors on lung fibroblasts in vitro. Int Arch Allergy Immunol 132: 168–176.

13. Takayama, G., K. Arima, T. Kanaji, S. Toda, H. Tanaka, S. Shoji, A. N. McKenzie, H. Nagai, T. Hotokebuchi, and K. Izuhara. 2006. Periostin: a novel component of subepithelial fibrosis of bronchial asthma downstream of IL-4 and IL-13 signals. The Journal of allergy and clinical immunology 118: 98–104.

14. Moynihan, B. J., B. Tolloczko, S. El Bassam, P. Ferraro, M. C. Michoud, J. G. Martin, and S. Laberge. 2008. IFN-gamma, IL-4 and IL-13 modulate responsiveness of human airway smooth muscle cells to IL-13. Respir Res 9: 84.

15. Punnonen, J., G. Aversa, B. G. Cocks, A. N. McKenzie, S. Menon, G. Zurawski, R. de Waal Malefyt, and J. E. de Vries. 1993. Interleukin 13 induces interleukin 4-independent IgG4 and IgE synthesis and CD23 expression by human B cells. Proc Natl Acad Sci U S A 90: 3730–3734.

16. Galli, S. J., M. Grimbaldeston, and M. Tsai. 2008. Immunomodulatory mast cells: negative, as well as positive, regulators of immunity. Nature reviews. Immunology 8: 478–486.

17. Galli, S. J., and M. Tsai. 2012. IgE and mast cells in allergic disease. Nature medicine 18: 693–704.

18. Toru, H., R. Pawankar, C. Ra, J. Yata, and T. Nakahata. 1998. Human mast cells produce IL-13 by high-affinity IgE receptor cross-linking: enhanced IL-13 production by IL-4-primed human mast cells. The Journal of allergy and clinical immunology 102: 491–502.

19. Chen, C. C., M. A. Grimbaldeston, M. Tsai, I. L. Weissman, and S. J. Galli. 2005. Identification of mast cell progenitors in adult mice. Proc Natl Acad Sci U S A 102: 11408–11413.

20. Moritz, D. R., H. R. Rodewald, J. Gheyselinck, and R. Klemenz. 1998. The IL-1 receptor-related T1 antigen is expressed on immature and mature mast cells and on fetal blood mast cell progenitors. J Immunol 161: 4866–4874.

21. Moussion, C., N. Ortega, and J. P. Girard. 2008. The IL-1-like cytokine IL-33 is constitutively expressed in the nucleus of endothelial cells and epithelial cells in vivo: a novel ‘alarmin’? PloS one 3: e3331.

22. Byers, D. E., J. Alexander-Brett, A. C. Patel, E. Agapov, G. Dang-Vu, X. Jin, K. Wu, Y. You, Y. Alevy, J. P. Girard, T. S. Stappenbeck, G. A. Patterson, R. A. Pierce, S. L. Brody, and M. J. Holtzman. 2013. Long-term IL-33-producing epithelial progenitor cells in chronic obstructive lung disease. The Journal of clinical investigation 123: 3967–3982.

23. Allakhverdi, Z., D. E. Smith, M. R. Comeau, and G. Delespesse. 2007. Cutting edge: The ST2 ligand IL-33 potently activates and drives maturation of human mast cells. J Immunol 179: 2051–2054.

24. Zheng, W., and R. A. Flavell. 1997. The transcription factor GATA-3 is necessary and sufficient for Th2 cytokine gene expression in CD4 T cells. Cell 89: 587–596.

25. Lavenu-Bombled, C., C. D. Trainor, I. Makeh, P. H. Romeo, and I. Max-Audit. 2002. Interleukin-13 gene expression is regulated by GATA-3 in T cells: role of a critical association of a GATA and two GATG motifs. J Biol Chem 277: 18313–18321.

26. Takemoto, N., Y. Kamogawa, H. Jun Lee, H. Kurata, K. I. Arai, A. O’Garra, N. Arai, and S. Miyatake. 2000. Cutting edge: chromatin remodeling at the IL-4/IL-13 intergenic regulatory region for Th2-specific cytokine gene cluster. J Immunol 165: 6687–6691.

27. Kishikawa, H., J. Sun, A. Choi, S. C. Miaw, and I. C. Ho. 2001. The cell type-specific expression of the murine IL-13 gene is regulated by GATA-3. J Immunol 167: 4414–4420.

28. Yao, X., W. Zha, W. Song, H. He, M. Huang, E. Jazrawi, P. Lavender, P. J. Barnes, I. M. Adcock, and A. L. Durham. 2012. Coordinated regulation of IL-4 and IL-13 expression in human T cells: 3C analysis for DNA looping. Biochem Biophys Res Commun 417: 996–1001.

29. Yao, X., Y. Yang, H. Y. He, M. Wang, K. S. Yin, and M. Huang. 2010. GATA3 siRNA inhibits the binding of NFAT1 to interleukin-13 promoter in human T cells. Chin Med J (Engl) 123: 739–744.

30. Kozuka, T., M. Sugita, S. Shetzline, A. M. Gewirtz, and Y. Nakata. 2011. c-Myb and GATA-3 cooperatively regulate IL-13 expression via conserved GATA-3 response element and recruit mixed lineage leukemia (MLL) for histone modification of the IL-13 locus. J Immunol 187: 5974–5982.

31. Monticelli, S., D. C. Solymar, and A. Rao. 2004. Role of NFAT proteins in IL13 gene transcription in mast cells. J Biol Chem 279: 36210–36218.

32. Masuda, A., Y. Yoshikai, H. Kume, and T. Matsuguchi. 2004. The interaction between GATA proteins and activator protein-1 promotes the transcription of IL-13 in mast cells. J Immunol 173: 5564–5573.

33. Li, B., M. R. Power, and T. J. Lin. 2006. De novo synthesis of early growth response factor-1 is required for the full responsiveness of mast cells to produce TNF and IL-13 by IgE and antigen stimulation. Blood 107: 2814–2820.

34. Li, B., J. Berman, P. Wu, F. Liu, J. T. Tang, and T. J. Lin. 2008. The early growth response factor-1 contributes to interleukin-13 production by mast cells in response to stem cell factor stimulation. J Immunotoxicol 5: 163–171.

35. Calo, E., and J. Wysocka. 2013. Modification of enhancer chromatin: what, how, and why? Mol Cell 49: 825–837.

36. Creyghton, M. P., A. W. Cheng, G. G. Welstead, T. Kooistra, B. W. Carey, E. J. Steine, J. Hanna, M. A. Lodato, G. M. Frampton, P. A. Sharp, L. A. Boyer, R. A. Young, and R. Jaenisch. 2010. Histone H3K27ac separates active from poised enhancers and predicts developmental state. Proc Natl Acad Sci U S A 107: 21931–21936.

37. Heintzman, N. D., R. K. Stuart, G. Hon, Y. Fu, C. W. Ching, R. D. Hawkins, L. O. Barrera, S. Van Calcar, C. Qu, K. A. Ching, W. Wang, Z. Weng, R. D. Green, G. E. Crawford, and B. Ren. 2007. Distinct and predictive chromatin signatures of transcriptional promoters and enhancers in the human genome. Nat Genet 39: 311–318.

38. Rada-Iglesias, A., R. Bajpai, T. Swigut, S. A. Brugmann, R. A. Flynn, and J. Wysocka. 2011. A unique chromatin signature uncovers early developmental enhancers in humans. Nature 470: 279–283.

39. Buenrostro, J. D., P. G. Giresi, L. C. Zaba, H. Y. Chang, and W. J. Greenleaf. 2013. Transposition of native chromatin for fast and sensitive epigenomic profiling of open chromatin, DNA-binding proteins and nucleosome position. Nat Methods 10: 1213–1218.

40. Corces, M. R., A. E. Trevino, E. G. Hamilton, P. G. Greenside, N. A. Sinnott-Armstrong, S. Vesuna, A. T. Satpathy, A. J. Rubin, K. S. Montine, B. Wu, A. Kathiria, S. W. Cho, M. R. Mumbach, A. C. Carter, M. Kasowski, L. A. Orloff, V. I. Risca, A. Kundaje, P. A. Khavari, T. J. Montine, W. J. Greenleaf, and H. Y. Chang. 2017. An improved ATAC-seq protocol reduces background and enables interrogation of frozen tissues. Nat Methods 14: 959–962.

41. Pierce, J. H., P. P. Di Fiore, S. A. Aaronson, M. Potter, J. Pumphrey, A. Scott, and J. N. Ihle. 1985. Neoplastic transformation of mast cells by Abelson-MuLV: abrogation of IL-3 dependence by a nonautocrine mechanism. Cell 41: 685–693.

42. Li, Y., B. Liu, L. Harmacek, Z. Long, J. Liang, K. Lukin, S. M. Leach, B. O’Connor, A. N. Gerber, J. Hagman, A. Roers, F. D. Finkelman, and H. Huang. 2018. The transcription factors GATA2 and microphthalmia-associated transcription factor regulate Hdc gene expression in mast cells and are required for IgE/mast cell-mediated anaphylaxis. J Allergy Clin Immunol 142: 1173–1184.

43. Bryksin, A. V., and I. Matsumura. 2010. Overlap extension PCR cloning: a simple and reliable way to create recombinant plasmids. Biotechniques 48: 463–465.

44. Tan, G., and B. Lenhard. 2016. TFBSTools: an R/bioconductor package for transcription factor binding site analysis. Bioinformatics (Oxford, England) 32: 1555–1556.

45. Spitz, F., and E. E. Furlong. 2012. Transcription factors: from enhancer binding to developmental control. Nature reviews. Genetics 13: 613–626.

46. Li, Y., X. Qi, B. Liu, and H. Huang. 2015. The STAT5-GATA2 pathway is critical in basophil and mast cell differentiation and maintenance. J Immunol 194: 4328–4338.

47. Wu, Z., A. J. Macneil, R. Junkins, B. Li, J. N. Berman, and T. J. Lin. 2013. Mast cell FcepsilonRI-induced early growth response 2 regulates CC chemokine ligand 1-dependent CD4+ T cell migration. J Immunol 190: 4500–4507.

