## Supplemental Table 1 for "The Clusters of Transcription Factors NFATC2, STAT5, GATA2, AP1, RUNX1 and EGR2 Binding Sites at the Induced *Il13* Enhancers Mediate *Il13* Gene Transcription in Response to Antigenic Stimulation"

**TableS1**

P75FP AAATAGATCTTGCCCAACAAAGCAGAGACC

P-1.0RP AATAAAGCTTGGCCTTAGCCTGTTGAAGC

P150FP AAATAGATCTTTAGCGGCCACTGGATTTT

P150_ALL MUT_FP AAATAGATCTTTAGCGGCCACTGGATGTCAAGTCAAAGAAAAAAAAAGTCAAGGTCAAGAGGTCGAGTCAATTTTCAGATAATGCCCAACAAAGC

P150_MUT_GAR_FP AAATAGATCTTTAGCGGCCACTGGATTTTCCAAAAAAGAAAAAAAAAAATTCAGTCAAGAGGTCGAGTCAATTTTCAGATAATGCCCAACAAAGC

P150_MUT_NS_FP AAATAGATCTTTAGCGGCCACTGGATGTCAAGTCAAAGAAAAAAAAAAATTCAAGATGAGTAAAGATG

P150_MUT_GA_FP AAATAGATCTTTAGCGGCCACTGGATTTTCCAAAAAAGAAAAAAAAAAATTCAGTCAAGAGGTCGATGTGGTTTTCAGATAATGCCCCAA

P150_MUT_NFAT_FP AAATAGATCTTTAGCGGCCACTGGATGTCAAAAAAAAGAAAAAAAAAAATTCAAGATGAGTAAAGATG

P150_MUT_STAT5_FP AAATAGATCTTTAGCGGCCACTGGATTTTCCGTCAAAGAAAAAAAAAAATTCAAGATGAGTAAAGATG

P150_MUT_HOXA5_FP AAATAGATCTTTAGCGGCCACTGGATTTTCCAAAAAAGAAAAAAAAAGTCAAGAGATGAGTAAAGATGTGGTTTTCAGATAA

P150_MUT_GATA2_FP AAATAGATCTTTAGCGGCCACTGGATTTTCCAAAAAAGAAAAAAAAAAATTCAGTCAAGGTAAAGATGTGGTTTTCAGATAATGCCCC

P150_MUT_AP1_FP AAATAGATCTTTAGCGGCCACTGGATTTTCCAAAAAAGAAAAAAAAAAATTCAAGGTCAAGGTCGATGTGGTTTTCAGATAATGCCCCAA

P150_MUT_RUNX1_FP AAATAGATCTTTAGCGGCCACTGGATTTTCCAAAAAAGAAAAAAAAAAATTCAAGATGAGTAAAGAGTCAATTTTCAGATAATGCCCAACAAAGC

E+8.9FP AAATGCTAGCTGATCATGGTTCCTTATCTGGACC

E+8.9RP AAATCTCGAGTGGCATTTGGCACATTAGAAAAAAATATGTAAG

E+6.5(+6534)FP AATAGAGCTCAAGGAGGTCTCTCTTCCAGTCC

E+6.5(+8153)RP AAATGCTAGCAGAGAAACCCTGTCTCGAAAACAAAAA

E+6.5_6534FP AATAGAGCTCAAGGAGGTCTCTCTTCCAGTCC

E+6.5_6770FP AATAGAGCTCAAGCCGCTAAAGCTGACAGCAA

E+6.5_6970FP AATAGAGCTCCTGAGAGCCTGTGGTGAGTG

E+6.5_7170FP AATAGAGCTCAAACACAGTCACTTCAGGGGTTTC

E+6.5_7370FP AATAGAGCTCTCCTCTGAGATGGTGGCAGG

E+6.5_7570FP AATAGAGCTCCCACCCAAAAGCGACAGAGC

E+6.5_7770FP AATAGAGCTCGCCTAAGAAACTCCCCCAGC

E+6.5_7970FP AATAGAGCTCTGATTCCTGGCTGATTCCTAGCC

E+6.5_7370FP AATAGAGCTCTCCTCTGAGATGGTGGCAGG

E+6.5_7420FP AATAGAGCTCTCCTGACCATGGGGAACCCC

E+6.5_7470FP AATAGAGCTCGCCTAGAGCTGTGTCCATAGC

E+6.5_7520FP AATAGAGCTCGTGAAAACTAAAAGTCACGAGGCCTC

E+6.5_570RP AAAT CTCGAG GCCGAGAAATGAATGAAGA

E+6.5_7470-570_MUT_STREFP AATAGGTACCGCCTAGAGCTGTGTCCATAGCCACCTTCCCCCACAGTGAGAACGGCGAAGGTGA

E+6.5_7470-570_MUT_EGR2FP AATAGGTACCGCCTAGAGCTGTGTCCATAGCAGAACGGCGAAGAACTGCCC

E+6.5_7470-570_MUT_FOXA2FP AATAGGTACCGCCTAGAAGAACGGCGAAGGCCACCTTCCCCCA

E+6.5_7470-570_MUT_EFSFP AATAGGTACCGCCTAGAAGAACGGCGAAGGCAGAACGGCGAAGAACTGAGAACGGCGAAGGTGAAAA

E+2.5FPAATGGTACCGTTTCACAGGTCTCCAGCC

E+2.5RP AAATCTCGAGAAACAACAACAAAAGCAAAAAACCAACA

E+0.2FP AATAGGTACCAAGAGGTCATGAGCAGGCT

E+0.2RP AAATCTCGAGAGACCGTGAGTAGACCGGTAG

E-1.0FP AAATGCTAGCAGGAGAGAGGTGAGGAGGA

E-1.0RP AAATCTCGAGTGCCTATTGGGTAGTGATGTCAAG

E-1.0 _E+0.2FP AAATGCTAGCCTGGCCTGAGGGAGGAAGTGGGTGAGGGTGCATCTTGACATCACTACCCAATAGGCAAAGAGGTCATGAGCAGGCTGG

E+0.2RP AGACCGTGAGTAGACCGGTAG

E+6.5_E+0.2OLFP_CTACCGGTCTACTCACGGTCTTCCTCTGAGATGGTGGCAGG

E+6.5_FP AAATGCTAGCTCCTCTGAGATGGTGGCAGG

E+6.5_RP AGCCGAGAAATGAATGAAGA

P150_OLFP TCTTCATTCATTTCTCGGCTCCTTTAGCGGCCACTGGATTTT

P75_E+0.2OLFP CTACCGGTCTACTCACGGTCTTGCCCAACAAAGCAGAGACC

P-1.0BglIIRP AAATAGATCTGGCCTTAGCCTGTTGAAGC
